# Bite force performance from wild derived mice has undetectable heritability despite having heritable morphological components

**DOI:** 10.1101/2021.12.03.470898

**Authors:** Samuel Ginot, Benedikt Hallgrímsson, Sylvie Agret, Julien Claude

## Abstract

Fitness-related traits tend to have low heritabilities. Conversely, morphology tends to be highly heritable. Yet, many fitness-related performance traits such as running speed or bite force depend critically on morphology. Craniofacial morphology correlates with bite performance in several groups including rodents. However, within species, this relationship is less clear, and the genetics of performance, morphology and function are rarely analyzed in combination. Here, we use a half-sib design in outbred wild-derived *Mus musculus* to study the morphology-bite force relationship and determine whether there is additive genetic (co-)variance for these traits. Results suggest that bite force has undetectable additive genetic variance and heritability in this sample, while morphological traits related mechanically to bite force exhibit varying levels of heritability. The most heritable traits include the length of the mandible which relates to bite force. Despite its correlation with morphology, realized bite force was not heritable, which suggests it is less responsive to selection in comparison to its morphological determinants. We explain this paradox with a non-additive, many-to-one mapping hypothesis of heritable change in complex traits. We furthermore propose that performance traits could evolve if pleiotropic relationships among the determining traits are modified.

## Introduction

It is generally assumed that the morphology of the musculoskeletal systems evolves in response to selection on performance (Herrel et al. 2005, Van Daele et al. 2008, Becerra et al. 2011, Ginot et al. 2017, Aerts et al. 2000, Vanhooydonck & Van Damme 2001). For this to occur, morphological variation must covary with performance (Arnold, 1983). For appendicular traits, relevant measures of performance might be speed (Blumstein et al. 2010, Zamora et al. 2014), endurance (Vanhooydonck et al. 2001) or range and durations of positional behaviors (Bezanson, 2017). For the morphology of the face and, in particular, the rodent skull, bite force is the performance measure most commonly thought to be relevant to fitness. Among species of mammals, bite force covaries with craniofacial morphology (Aguirre et al. 2002, Herrel et al. 2008, Freeman & Lemen 2008, Van Daele et al. 2008, Becerra et al. 2011), presumably due, at least in part, to the morphological response to selection on this key performance parameter. Within species, however, the relationship between craniofacial morphology and bite force performance is often less clear (e.g. Herrel et al. 2005, Van Daele et al. 2008, Becerra et al. 2011, Ginot et al. 2017) and even more so within age-cohorts of a population (Van Daele et al. 2008, Becerra et al. 2011, Ginot et al. 2017, 2020). These results suggest that bite force is a multi-factorial trait, and that intraspecific variation in bite force is not only due to variation in morphology, but also depends on behavioural or environmental variation.

For performance traits such as bite force to evolve, they must be heritable. Quantitative genetics studies in the wild or in controlled conditions tend to show that morphological traits are more heritable than fitness-related traits (e.g. life-history, behavioural, performance traits ; Mousseau & Roff 1987, Houle 1992, Hoffman et al. 2016). The low heritability of fitness-related traits is often explained through Fisher’s (1958) theorem of natural selection, which states that fitness-related traits will have lower additive genetic variance because they are under stronger stabilizing selection than other traits (Mousseau & Roff 1987), reducing genetic variation, and possibly increasing canalization, to ensure sufficient performance for survival. However, Houle (1992) also proposed that low heritabilities in fitness-related traits might be due to larger amounts of non-additive genetic and environmental variation compared to morphological traits. These two non-exclusive views have different evolutionary implications because a trait with low additive genetic variance (even with low non-additive and environmental variance) will be less easily evolvable than one with high additive genetic variance (Houle 1992, Hoffman et al. 2016). Therefore, Houle (1992) proposed that quantitative genetics studies should always report the partitioning of variance of a trait between the additive genetic and other effects rather than to report only the heritability.

Numerous studies have documented links between morphological variation and performance (endurance, speed, bite force) with the aim of better understanding the functional relationship between phenotype and fitness (e.g. Herrel et al. 2005, Van Daele et al. 2008, Becerra et al. 2011, Ginot et al. 2017, Aerts et al. 2000, Vanhooydonck & Van Damme 2001). However, these studies rarely include quantitative genetic analyses because it requires phenotyping both morphological traits and performance within a controlled pedigree context. These studies are practically non existent for wild or wild derived populations of mammals, with the recent exception of a study on skull morphology and bite force in mouse lemurs (Zablocki-Thomas et al. 2021; see also Garland 1988, Tsuji et al. 1989, Noble et al. 2014 for studies of morphology and running performance heritability in lizards and snakes). Selection should act on performance, driving morphological change (Arnold 1983). Therefore, heritable variation and covariation between both is expected, a result supported by Zablocki-Thomas et aL (2021). Since morphology, function and performance are related, using quantitative genetic approaches may help reconcile their differential heritabilities, and better understand their co-evolution at the interspecific scale.

This study therefore has several objectives: i) identifying the morphological characters that relate to bite force in our population of lab mice; ii) quantifying the heritability and importance of the additive genetic, dominance, and environmental variation in *in vivo* bite force and skull morphology (size, shape and morpho-functional traits); iii) proposing hypotheses regarding the co-evolution of characters with different heritabilities by comparing intra and interspecific patterns of morphological variation. Based on previous studies and theory, we expect that i) skull size and cranio-mandibular morphology should be correlated with bite force, ii) bite force should be less heritable than morphological characters, and iii) that characters that commonly vary at a macroevolutionary level should correspond to the most heritable ones.

## Materials and Methods

### Specimens

All mice (*Mus musculus*) used in this study are from a colony of mice bred in the lab from wild ancestors captured on Mainland, the biggest of the Orkney Islands (Scotland). A half-sib design pedigree in which each father was bred with three different and unrelated females was produced. In total, we used 18 fathers (“sires”) that reproduced with 54 mothers (“dams”), and gave birth to 336 offspring. Litter sizes varied among families from 3 to 10 pups, with a mean of 6.2 pups. The number of specimens was limited by the wild origin of the colony, the need to cross genetically unrelated individuals, and the ethical necessities of raising vertebrate animals in refined conditions, and to reduce the number of specimens killed. Animals were treated in accordance with the guidelines of the American Society of Mammalogists, and within the European Union legislation guidelines (Directive 86/609/EEC and 2010/63/UE). All lab procedures were carried out under approval no. A34-172-042 (DDPP Hérault Prefecture).

### Bite force measurements

Bite force measurements were performed when the offspring were strictly 68 days old. All *in vivo* bite force data were recorded at the incisors using a Kistler force transducer linked to a charge amplifier, similar to the set-up presented in Herrel et al. (1999) and Aguirre et al. (2002). We performed three consecutive trials for each animal. The maximal bite force recorded across the three trials was retained and used in subsequent analyses. The mice were then euthanized by CO2 inhalation. We also measured the parents’ bite force in the same fashion, although their age varied.

### Morphometric data

Using the software TPSDig2.0 (Rohlf 2010), 24 landmarks were digitized on the crania of the parents and offsprings in palatal view, and 16 landmarks on the mandibles in lateral jugal view (Fig. 1A-B). All coordinates data were imported in R (R Core Team 2017) and the shapes were centered and superimposed using functions from Claude (2008). We also computed centroid sizes for the mandibles and crania.

**Figure 1.**
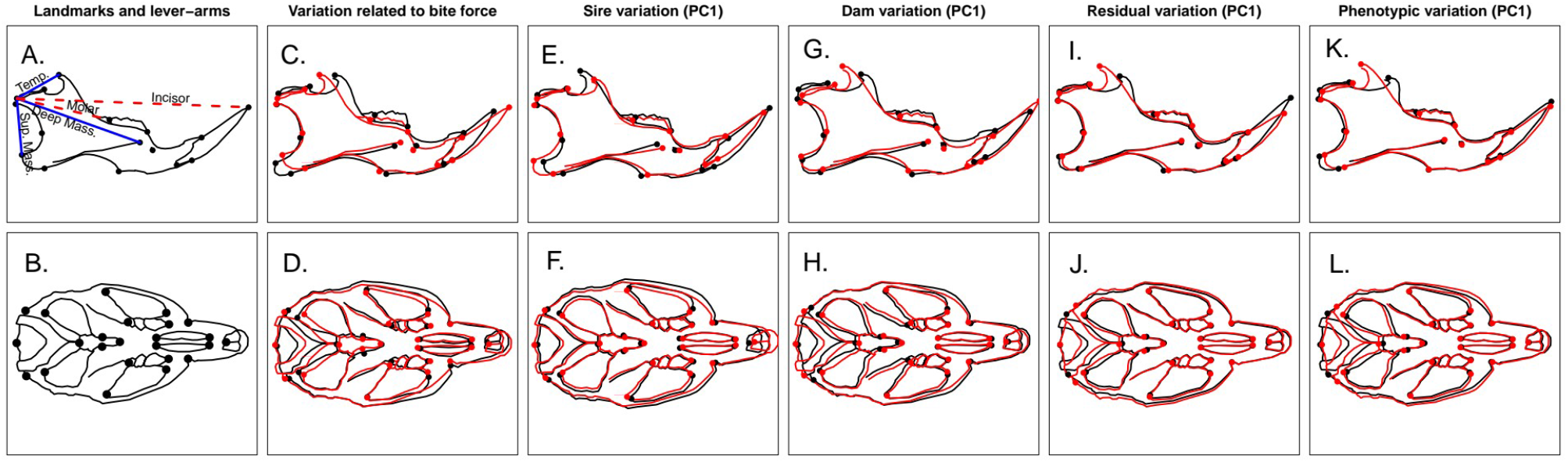
**A**. Mouse mandible outline in lateral view showing homologous landmarks and linear measurements corresponding to the lever-arms studied here. Solid blue lines represent in-levers for the temporal (Temp.), deep masseter (Deep Mass.), and superficial masseter (Sup. Mass.) ; dashed red lines represent out-levers at the incisor or at the molar. **B**. Mouse cranium outline in palatal view showing homologous landmarks used in this study. **C-D**. Patterns of cranial and mandibular shape variation related to bite force obtained by projecting measured bite forces on a multivariate linear regression of shape on bite force. Black and red shapes respectively represent maximal and minimal deformations, amplified five times for better visualization. **E-L**. Patterns of cranial and mandibular shape variation. Black and red shapes respectively represent maximal and minimal deformations for the first principal component of variation, but note that the orientation of the axes are arbitrary and therefore not necessarily the same across components. Sire variation corresponds to the heritable component of phenotypic variation. Dam variation corresponds to the maternal (i.e. dominance and common nest) and heritable component of variation. Residual variation corresponds to the residual individual variation including environmental variation. Phenotypic variation corresponds to the total shape variation.

In addition to these geometric morphometric data, we also used our landmarks to calculate univariate functional distances on the mandible (Fig. 1B). These traits represent in-lever lengths for three of the adductor muscles (temporal, deep masseter, and superficial masseter), and out-lever lengths at the molar and incisor. Using these lengths, we calculated the mechanical advantage (MA) for the various levers for each individual as MA = In-lever length / Out-lever length (Radinsky 1981). Phenotypic correlation matrix were computed for these functional traits, weight, and bite force using Pearson’s product-moment correlation.

Bite force-related global shape variation was modelled using a multivariate regression of shape on bite force. Using the ‘predict’ function, modelled shapes were reconstructed, and individuals with the maximum and minimum bite force recorded were used to represent extreme cranium or mandible shape variation related to bite force. The differences being fairly small, they were amplified to be visible graphically.

### Quantitative genetic analyses

Using *in vivo* bite force and selected univariate morphometric measurements (centroid sizes, inlever and outlever lengths, mechanical advantages), we calculated one parent-offspring regressions for the mother and father separately. We also tested differences between male and female offsprings using Welch’s t tests, and within the linear regression models by using sex of the offspring as an explanatory variable. Mother-offspring regressions tend to overestimate heritabilities because they include maternal effects, which is not the case with father-offspring regressions. This allowed us to get a first estimate of heritability (h^2^ = 2^*^slope; Falconer 1989), with associated 95% confidence intervals computed from the standard error and degrees of freedom for the slope. Parent-offspring regressions suffer from several biases (notably because the parents are of variable ages) and do not allow to easily partition the variance between additive genetic (V_A_) *versus* other effects. To achieve this, we used a mixed effect model, with the ‘mmer’ function from package sommer in R (Covarrubias-Pazaran 2016). For each trait (centered but not scaled) we computed mixed models of the trait with sire and dam as random effects. This allowed us to partition the phenotypic variance of our traits into a sire component (σ^2^_sires_ = 1/4*V_A_), a dam component (σ^2^_dams_ = 1/4*V_A_ + 1/4*V_D_ + V_EC_), and residual component (σ^2^_residual_ = 1/2*V_A_ + 3/4*V_D_ + V_EW_). V_A_ is the additive genetic variance, V_D_ the dominance variance, V_EC_ being the common evironmental variance of full sibs and V_EW_ the environmental variance within a litter. We then calculated heritability as follows:

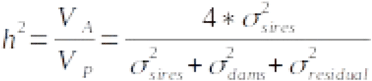 (Falconer 1989). For additive genetic variance estimates (mmer mixed models), we computed confidence intervals by jackknifing sires of our sample and taking the 0.975 and 0.025 quantiles of the computed pseudo-values. These quantile values were then passed into the heritability formula to obtain confidence intervals for mixed model heritability. This allowed us to assess whether the number of sires used in the study was large enough allowed us to actually obtain heritability estimates with some degree of confidence. If confidence intervals for heritabilities all included 0, then our sample clearly would not allow us to detect any heritable variation. We also calculated the evolvability of the same traits as: 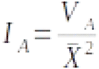 (Houle 1992), where X bar is the mean of the trait studied.

### Variance-covariance and correlation matrices between traits

Still using the ‘mmer’ function, we computed sire, dam and residual covariance estimates between pairs of traits, with two traits as the dependent variables and again sire and dam as random factors. We combined these with the variances previously computed to obtain variance-covariance matrices for the chosen linear traits and for cranium and mandible shape. Total phenotypic, additive genetic (sire), dam and residual components of mandible shape, cranium shape and morpho-functional traits were thereby obtained. When possible (i.e. when trait additive genetic variance were significantly different from 0), correlation matrices were computed from the variance-covariance matrices. For the sire variance-covariance matrix, many correlations between pairs of traits were outside the [-1,1] range, due to error in variance of covariance estimates which sometimes produced covariance larger than standard deviation products. Based on the variance-covariance matrices for shape, we computed a multivariate heritability estimate, following Monteiro et al. (2002). This index consists in dividing the sum of the diagonal values of the additive genetic variance-covariance matrix by the sum of the diagonal of the total phenotypic variance-covariance matrix, and dividing by 0.25 for half-sib designs. This index is given for providing a general idea of shape heritability but we are, aware of potential biases as Klingenberg and Monteiro (2005) have raised issues with this estimate when matrices are not isotropic; something that was likely the case here due to unsolvability of some of our variance-covariance matrices. In consequence, results for multivariate heritability and correlations between matrices are reported as Supplementary Information for disclosure, but should be taken cautiously. Finally, we graphically represented the first principal components of shape variation by projecting the original superimposed coordinates on the variance-covariance matrices (sire, dam, residual and phenotypic) to identify whether major axes of additive genetic (sire) and other variation might relate with morphological correlates of bite force.

## Results

### Phenotypic correlations between bite force and morphology

Bite force is correlated with centroid size as well as shape of both the mandible and the cranium (Fig. 1, SI Fig. 1). Higher bite force is related to more anteriorly positioned coronoid process (temporal muscular insertion), more posteriorly developed angular process (superficial masseter and pterygoid muscular insertion), shorter and higher mandible (i.e. more ‘robust’ morphologies) (Fig. 1C). In the cranium, variation related to bite force was less conspicuous, with higher bite force related to slightly shorter and wider crania with larger temporal arches and fossa (Fig. 1D). Some lever-arms and mechanical advantages of the mandible are also correlated with bite force (SI Fig. 1). These are the in-lever of the superficial masseter (related to posteriorly and ventrally positioned angular process), the mechanical advantages for the superficial masseter/incisor, the temporal/incisor (related to the position of coronoid process), and the temporal/molar levers, and negatively related to the incisor out-lever (e.g. length of the mandible from the condylar process to the tip of incisors, Fig. 1, SI Fig. 1).

From these results, a subset of univariate traits were chosen for which we obtained additive genetic variance, heritability estimates and genetic correlation when it was possible: bite force, centroid size (cranium and mandible), superficial masseter in-lever length, superficial masseter/incisor mechanical advantage, temporal/incisor mechanical advantage, and out-lever lengths.

### Parent-offspring regressions

Parent-offspring regressions for the selected characters (Fig. 2) revealed differences between father and mother regressions (Table 1). Father-offspring regressions suggested significant (p < 0.05) heritable variation for cranium and mandible centroid size (h^2^ = 0.25 and h^2^ = 0.43 respectively, Fig. 2). The heritability of the temporal/incisor out-lever length mechanical advantage is 0.4, 0.67 for the incisor out-lever length mechanical advantage and 0.51 for the molar out-lever length mechanical advantage (Fig. 2, Table 1). However, the slopes were not significant for bite force and for the superficial masseter in-lever and mechanical advantage and heritabilities were therefore low and not quantifiable with significance for these variables (Table 1). Mother-offspring regressions were significant for all characters except for the superficial masseter in-lever length. When slopes were significant, the differences between the slopes for male and female offspring were not significant (all p > 0.1, Table 1) despite the intercept being generally significantly different (p < 0.01, i.e. significant sexual dimorphism, see SI Fig. 2; Table 1). The incisor out-lever length appeared to show different slopes for males and females (Fig. 2), but even in this case, the difference was not significant (Estimate = -0.225, S.E. = 0.139, t = -1.625, df = 330, p > 0.1). This showed that, although characters were sexually dimorphic (i.e. regressions had different intercepts for males and females, Table 1, SI Fig. 2), the patterns of transmission of variability were not different between sexes (i.e. the slopes were not different). Therefore, we pooled males and females in the following mixed model analyses, and added sex as a fixed effect in the cases where univariate variables were sexually dimorphic (Fig. 2, SI Fig. 2).

**Figure 2.**
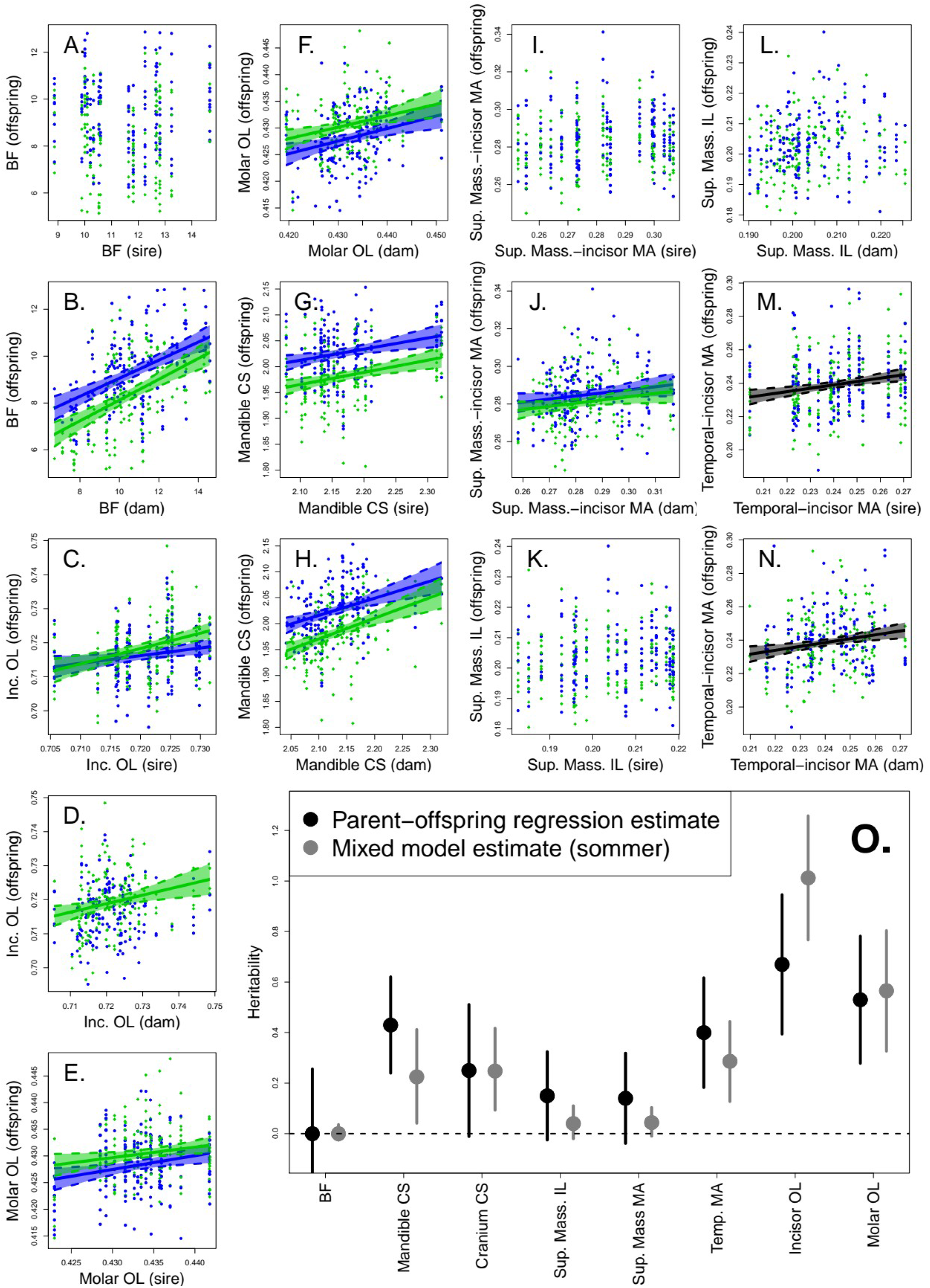
**A-N**. Parent-offspring regressions for chosen univariate morphometric traits and bite force. Blue circles are male offspring, green diamonds are female offspring. Significant regression lines were plotted separately in green and blue when a significant sex effect was detected, or in black when no significant difference between males and females was found. Dashed lines and transparent areas represent 95% confidence intervals on the regression line. **O**. Heritability estimates for univariate morphometric traits and bite force, obtained from the slope of parent-offspring regression or mixed models. Whiskers represent 95% confidence intervals for the heritability estimates. BF: Bite force; CS: Centroid size; Inc.: Incisor; IL: In-lever; MA: Mechanical advantage; OL: Out-lever; Sup. Mass.: Superficial masseter.

**Table 1.** Results of the parent-offspring regressions for bite force, size and morphometric traits in our pedigree of wild-derived *Mus musculus*.

|  | Estimate | Std.Error | t value | Pr(>\|t\|) |
| --- | --- | --- | --- | --- |
| **BF offspring ~ BF dam \* sex offspring)** |  |  |  |  |
| (Intercept) | 13.77563 | 1.15076 | 11.971 | <2e-16*** |
| BF dam | 0.25937 | 0.04624 | 5.609 | 4.4e-08*** |
| Sex offspring | 3.23767 | 1.65297 | 1.959 | 0.051. |
| BF dam : Sex offspring | -0.04261 | 0.06567 | -0.649 | 0.517 |
| --- |  |  |  |  |
| **BF offspring ~ BF sire \* sex offspring)** |  |  |  |  |
| (Intercept) | 20.220662 | 1.224866 | 16.51 | <2e-16*** |
| BF sire | -0.006004 | 0.046295 | -0.13 | 0.8969 |
| Sex offspring | 3.144052 | 1.786026 | 1.76 | 0.0793. |
| BF sire : Sex offspring | -0.030707 | 0.066689 | -0.46 | 0.6455 |
| --- |  |  |  |  |
| **Out-lever incisor offspring ~ Out-lever incisor sire \* sex offspring** |  |  |  |  |
| (Intercept) | 0.3931 | 0.0727 | 5.407 | 1.23e-07*** |
| Out-lever incisor sire | 0.4517 | 0.1008 | 4.483 | 1.02e-05*** |
| Sex offspring | 0.1602 | 0.1001 | 1.601 | 0.110 |
| Out-lever incisor sire : Sex offspring | -0.2254 | 0.1387 | -1.625 | 0.105 |
| --- |  |  |  |  |
| **Out-lever incisor offspring ~ Out-lever incisor dam \* sex offspring** |  |  |  |  |
| (Intercept) | 0.53846 | 0.05806 | 9.275 | <2e-16*** |
| Out-lever incisor dam | 0.25051 | 0.08045 | 3.114 | 0.00202** |
| Sex offspring | 0.1116 | 0.08218 | 1.358 | 0.17541 |
| Out-lever incisor dam : Sex offspring | -0.15837 | 0.11391 | -1.390 | 0.16540 |
| --- |  |  |  |  |
| **Centroid size offspring ~ Centroid size dam \* sex offspring** |  |  |  |  |
| (Intercept) | 1.13019 | 0.15784 | 7.16 | 5.68e-12*** |
| Dam size | 0.40019 | 0.07403 | 5.405 | 1.28e-07*** |
| Sex offspring | 0.18956 | 0.22679 | 0.836 | 0.404 |
| Dam size : Sex offspring | -0.06865 | 0.10646 | -0.645 | 0.52 |
| --- |  |  |  |  |
| **Centroid size offspring ~ Centroid size sire \* sex offspring** |  |  |  |  |
| (Intercept) | 1.47232 | 0.12687 | 11.605 | <2e-16*** |
| Sire size | 0.23483 | 0.05837 | 4.024 | 7.11e-05*** |
| Sex offspring | 0.08816 | 0.19609 | 0.45 | 0.653 |
| Sire size : Sex offspring | -0.01965 | 0.0903 | -0.218 | 0.828 |
| --- |  |  |  |  |
| **In-lever MS offspring ~ In-lever MS sire \* sex offspring** |  |  |  |  |
| (Intercept) | 0.191029 | 0.012363 | 15.452 | <2e-16*** |
| In-lever MS sire | 0.053393 | 0.061022 | 0.875 | 0.382 |
| Sex offspring | -0.001221 | 0.018226 | -0.067 | 0.947 |
| In-lever MS sire : sex offspring | 0.016989 | 0.089296 | 0.190 | 0.849 |
| --- |  |  |  |  |
| **In-lever MS offspring ~ In-lever MS dam \* sex offspring** |  |  |  |  |
| (Intercept) | 0.172328 | 0.018211 | 9.463 | <2e-16*** |
| In-lever MS dam | 0.145298 | 0.089329 | 1.627 | 0.105 |
| Sex offspring | -0.001477 | 0.025153 | -0.059 | 0.953 |
| In-lever MS dam : sex offspring | 0.016980 | 0.123147 | 0.138 | 0.890 |
| --- |  |  |  |  |
| **Mechanical advantage MS/inc offspring ~ Mechanical advantage MS/inc sire \* sex offspring** |  |  |  |  |
| (Intercept) | 0.266857 | 0.017531 | 15.222 | <2e-16*** |
| Mechanical advantage sire | 0.049657 | 0.062403 | 0.796 | 0.427 |
| Sex offspring | -0.004298 | 0.025717 | -0.167 | 0.867 |
| Mechanical advantage sire : sex offspring | 0.029765 | 0.0985 | 0.328 | 0.743 |
| --- |  |  |  |  |
| **Mechanical advantage MS/inc offspring ~ Mechanical advantage MS/inc dam \* sex offspring** |  |  |  |  |
| (Intercept) | 0.232134 | 0.025315 | 9.170 | <2e-16*** |
| Mechanical advantage dam | 0.17242 | 0.089571 | 1.925 | 0.0551. |
| Sex offspring | 0.006261 | 0.034577 | 0.181 | 0.8564 |
| Mechanical advantage dam : sex offspring | -0.008645 | 0.12208 | -0.071 | 0.9436 |
| --- |  |  |  |  |
| **Mechanical advantage T/inc offspring ~ Mechanical advantage T/inc sire \* sex offspring** |  |  |  |  |
| (Intercept) | 0.18719 | 0.017691 | 10.581 | <2e-16*** |
| Mechanical advantage sire | 0.2125 | 0.073602 | 2.887 | 0.00414** |
| Sex offspring | 0.008273 | 0.026871 | 0.308 | 0.75835 |
| Mechanical advantage sire : sex offspring | -0.027717 | 0.11155 | -0.248 | 0.80392 |
| --- |  |  |  |  |
| **Mechanical advantage T/inc offspring ~ Mechanical advantage T/inc dam \* sex offspring** |  |  |  |  |
| (Intercept) | 0.16707 | 0.02357 | 7.089 | 8.88e-12*** |
| Mechanical advantage dam | 0.29587 | 0.09785 | 3.024 | 0.0027** |
| Sex offspring | 0.03278 | 0.03352 | 0.978 | 0.3289 |
| Mechanical advantage dam: sex offspring | -0.13218 | 0.13891 | -0.952 | 0.3420 |
| --- |  |  |  |  |
| **Mechanical advantage T/mol offspring ~ Mechanical advantage T/mol sire \* sex offspring** |  |  |  |  |
| (Intercept) | 0.30593 | 0.03002 | 10.193 | <2e-16*** |
| Mechanical advantage sire | 0.23043 | 0.07518 | 3.065 | 0.00236** |
| Sex offspring | 0.02757 | 0.045 | 0.613 | 0.54048 |
| Mechanical advantage sire : sex offspring | -0.0612 | 0.11232 | -0.545 | 0.58621 |
| --- |  |  |  |  |
| **Mechanical advantage T/mol offspring ~ Mechanical advantage T/mol dam \* sex offspring** |  |  |  |  |
| (Intercept) | 0.25574 | 0.04059 | 6.301 | 9.9e-10*** |
| Mechanical advantage dam | 0.35402 | 0.10087 | 3.51 | 0.000514**<br>* |
| Sex offspring | 0.0858 | 0.05762 | 1.489 | 0.137488 |
| Mechanical advantage dam: sex offspring | -0.20758 | 0.14319 | -1.450 | 0.148142 |

### Mixed effect model analyses

The mixed effect linear model analyses run on the selected traits allowed us to partition the phenotypic variation between sire variance (related to additive genetic variance(σ^2^_sires_ = 1/4*V_A_), and dam variance and residual variance (involving additive genetic, dominance, environmental and error variance; SI Table 1). Looking at the sire, dam and residual components of variance, it appears that bite force has the lowest sire variance, while having the highest dam and residual components (and generally highest variance overall). In comparison, all other variables have much lower total variance (therefore much lower dam and residual components). Among these traits, centroid sizes had both high sire and high dam and residual, followed by mechanical advantages (except for the superficial masseter, which had much lower sire variance than the temporal mechanical advantages). Although the in-levers and out-levers had rather low sire variance, they also had much lower dam and residual components than mechanical advantages or centroid sizes. In particular, the incisor and molar out-levers had much lower dam and residual components than all other variables.

Accordingly with the levels of additive genetic variance versus total phenotypic variance, heritability (Fig. 2O) for bite force is not detected (not significantly different from zero), but it is high for the out-lever lengths especially the incisor out-lever length (molar out-lever h^2^ = 0.55, incisor out-lever h^2^ = 0.94). Centroid sizes and the temporal/incisor mechanical advantage have intermediate heritabilities (cranium centroid size h^2^ = 0.17, mandible centroid size h^2^ = 0.16, mechanical advantage h^2^ = 0.29) with confidence intervals not including 0, while the superficial masseter in-lever and mechanical advantage have low heritabilities (h^2^ = 0.04 for both) which are not different from 0 according to confidence intervals. These results are similar to the estimates obtained from the father-offspring regressions although errors are larger for parent-offspring estimates (Fig. 2).

Evolvability of the traits shows divergent results from heritability (SI Table 1). However, since bite force Va is the lowest, bite force evolvability is also the lowest. Contrary to heritability, the out-levers have fairly low evolvability (especially the molar out-lever), as do centroid sizes and in-lever for the superficial masseter. Only the temporal mechanical advantage has a notably larger evolvability.

Sire, maternal and residual correlation matrices were computed from their corresponding variance-covariance matrices (SI Fig. 1). However, the sire component correlations between traits cannot be computed for all pairs of traits (notably for bite force which had no detectable additive genetic variance). These problems did not arise for the other components of variance-covariance and correlation. The variance-covariance patterns of shape change are illustrated in Fig. 1. It appears clearly that for the mandible, the sire (additive genetic) pattern of variation differs from the dam pattern and phenotypic pattern. Notably, the degree of anterior/posterior variation of the coronoid process (smaller for additive genetic effects), and anterior/posterior projection of the angular process (larger for the additive genetic effects). Dam and phenotypic patterns are similar, notably for the coronoid and angular process. Finally, the residual variation pattern is very similar to the dam and phenotypic pattern for the coronoid process, but displays a unique pattern of variation in the incisor, probably reflecting its growth. For the cranium, the main additive genetic variation was in the shorter snout and wider zygomatic arches, a pattern not observed in the dominance, environmental and phenotypic variation. Dam, residual and phenotypic patterns varied mostly in the posterior (occipital) region.

## Discussion

### Phenotypic bite force-morphology correlation

As expected, we determined that morphology and performance are linked at the intra-specific level. The strongest predictor of bite force is size, with a correlation coefficient of about 0.4 (SI Fig. 1), which could be explained by larger mandibles having larger and stronger muscles (Ginot et al. 2018). However, while morphology does correlate significantly with bite force (Fig. 1, SI Fig. 1), the correlation is relatively low suggesting that a large part of realized (i.e. measured) bite force performance variation is due to other factors (Ginot et al. 2017, 2020).

### Heritability of bite force and morphology

Parent-offspring regressions clearly show that realized *in vivo* bite force is not significantly heritable in our study (Fig. 2A, O), while elements of skull morphology that relate to jaw mechanics are (Fig. 2C-H, J, M-N, O). The mixed effect model analyses show that the absence of significant bite force heritability is not only due to high residual variance (Houle 1992), but also to the fact that additive genetic effects cannot be detected in the variance partitioning of this trait (SI Table 2). On the other hand, morphological characters display various levels of additive genetic variance. As expected, heritability and evolvability of the characters studied show divergent results (Houle 1992). Out-levers (related to mandible length), centroid sizes and temporal mechanical advantage have high heritabilities, while only temporal mechanical advantage is highly evolvable. However, both heritability and evolvability indices suggest that realized bite force may be unresponsive to selection, while morphological traits will show varying levels of response. It should be kept in mind, however, that the sample size of sires involved in this study may not be sufficient to detect low additive genetic variance and therefore heritability, as may be the case for bite force. Therefore, we obviously cannot affirm that bite force has absolutely no additive genetic component (proving an absence is logically impossible), but it is comparatively much lower than for morphological traits, for which we do detect additive genetic variance. Low signal to noise ratio is in any case expected in quantitative genetics studies (Falconer 1989). Our results appear more extreme, but similar to those reported by Zablocki-Thomas et al. (2021), who found low heritability for bite force compared to associated morphological traits. This study had the advantage of a larger sample size, building upon a colony of wild mouse lemurs which has been started from wild individuals over 50 years ago. Our study could not equal this sample size, but still had enough power to detect intermediate and high additive genetic variances, and may suffer less from biases due to evolution in captivity. It should also be noted that despite a larger sample size, Zablocki-Thomas and colleagues (2021) also reported non-significant weak heritability for another performance trait: pull strength, which demonstrates that, even within the same population, different performance traits may or may not be detectably heritable.

An important feature in our study is that the mice were reared in laboratory conditions, and so not subjected to extreme and varied diets. Laboratory conditions likely involve reduced environmental variance (reduced variation of food hardness) compared to natural conditions. While reduction in environmental variance would, in principle, increase heritability (by reducing the denominator of the heritability equation), it is possible that the opposite would actually occur for bite force: the reduced range of food hardness can limit the genetic expression of developmental variation related to food hardness variability. In other terms, genetic-environment (GxE) interactions are probably modified. More stressful environmental conditions might reveal genetic variation in bite force because individual animals use a larger range of their available bite force to process food of varying physical properties (Fig. 3A). A recent paper on wild-derived strains of *Peromyscus leucopus* (Lacy et al. 2018) confirms that heritability and addivitive genetic variance change through generations of captivity. The changes however were not consistent (both increases and decreases were found) across generations and across traits. Similarly, we do not know what would happen under conditions of directional selection created by a change in diet requiring an increase in bite force under natural conditions. Further studies, if possible in the wild or semi-captivity, are therefore needed to investigate how changing environmental conditions could alter the genetic variance-covariance among characters. That caveat aside, it is striking that despite the reduced environmental variance of laboratory conditions, the heritability of bite force is reduced to the point of undetectability.

**Figure 3.**
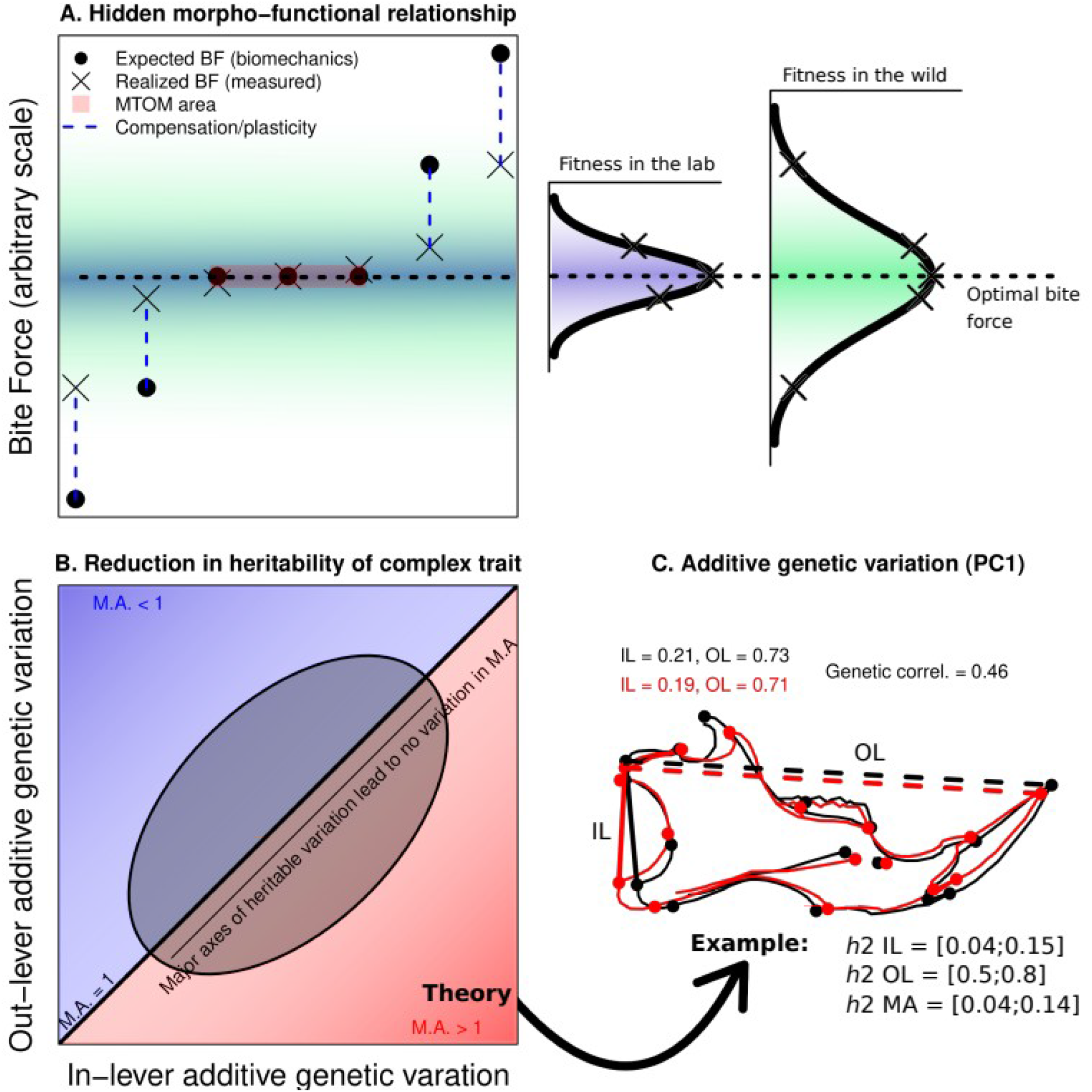
**A**. Theoretical graph illustrating several factors that can blur the relationship between morphology and performance, and reduce heritability. Each black dot represents an individual’s expected bite force based on its unique morphology (e.g. based on biomechanical model). The crosses represent the realized bite force, which can differ from the expected bite force due to plastic changes such as muscular compensation or behavioural differences (blue vertical dashed lines). In the example we consider that this plasticity tends to bring the realized bite force closer to the optimal bite force (dashed horizontal line), but also that this plasticity is limited for every indivividual. This plasticity will therefore reduce heritable variation in realized bite force, while at the same time blurring the morpho-functional relaptionship, so heritable morphological variation will be less visible in the realized bite force than it would in expected bite force. The three individuals in the middle represent the effect of many-to-one mapping: despite having different morphologies, their bite force performance will be the same, which again will blur the morpho-functional relationship. Finally, the fitness gradients related to bite force show that these effects may be even stronger in laboratory conditions: keeping animals in standard conditions with only one type of food, we may have selected out the extreme individuals, that could not compensate in terms of bite force, and less extreme individuals would tend to plastically respond to these lab conditions. In the wild, extreme individuals may have maintained thanks to more variable food sources, and other individuals could plastically reach higher fitness, or have more similar realized and expected bite forces. **B**. Additionally, we propose that complex traits, such as mechanical advantage (MA), which depend on two or more unidimensional traits, here the corresponding in-lever (IL) and out-lever (OL), may present reduced heritability. Indeed, these complex traits generally do not depend on additive relationships between their components, and frequently show many-to-one mapping. Here, MA is a ratio of IL over OL. The figure shows that, in the case where the IL and OL are both heritable and genetically positively correlated, the major axis of heritable covariation between them corresponds to no heritable variation in MA (MA ∼ 1, as shown by the ellipse and lighter gradient colors). **C**. This example illustrates that the mechanism suggested in B. does happen in real-life data. Here we show the superficial masseter IL (solid lines) and OL at the incisor (dashed lines). It appears that both are genetically correlated, as can be seen on the outline (black shape has longer IL and longer OL compared to red shape). Heritability of the superficial masseter MA is about 0, despite the OL being the most heritable trait found in this study. It should be noted that this effect not only reduces heritability, but more precisely reduces additive genetic variation, and could be extended in more complex traits such as performance or fitness.

Non-exclusive from the phenomena explained in the previous paragraph, the reduced additive genetic variance for bite force may also be due to strong stabilizing selection on this fitness-related character (i.e. Fisher’s theorem). Indeed, because mechanically expected bite force performance is related to the optimal ability to feed (Fig. 3A), its margin for variation is expected to be low (Cheverud 1996). Our study would be an extreme case but is consistent with both theoretical expectations and empirical studies (Fisher 1958, Mousseau & Roff 1987, Hoffman 2016). Due to its reduced additive genetic variance, bite force is genetically uncorrelated with morphology, despite covarying phenotypically. It would therefore appear that the phenotypic morphology-performance correlation emerges from sources other than the genetic correlation such as environmental variance or maternal factors. However, if bite force additive genetic variance had been detected, it may have been genetically correlated to morphological traits, as was found by Zablocki-Thomas et al. (2021).

### Disconnection between heritabilities of a complex trait and its components

The magnitude of additive genetic variance of a trait determines its evolvability (Fisher 1958, Houle 1992). Our results therefore suggest that selection on realized bite force in itself will resist more to evolutionary changes than morphology. The conundrum here is that bite force correlates with morphology, which is heritable, and yet bite force is not itself heritable (or much less heritable than morphology). This might occur because bite force is determined not only by bone but also by muscular structure. However, muscle size and morphology have a well known and strong relationships with bone morphology including the shape of the mandible and skull (Herring 1993). Further, muscular morphology is also heritable. An alternative explanation we propose is that bite force has a complex relationship to its morphological basis such that multiple and quite different anatomical arrangements generate similar bite forces (i.e. a many-to-one relationship). As shown in Figure 3, heritable variation in the in and out-levers of the mandible might translate to no heritable variation in bite force if additive genetic variances for these traits covary such as to produce a region over which the additive genetic variance for mechanical advantage is essentially invariant. Since both variables correlate imperfectly with bite force, this region is an ellipse rather than a line which also means that there is effectively a multivariate range of morphological variation that is compatible with genetically invariant bite force. Our data provides some support for this idea (Fig. 3C, SI Fig. 1). The in-lever (IL) and incisor out-lever (OL) length of the superficial masseter muscle both show some detectable additive genetic variance, and therefore both are heritable (SI Table 1), the out-lever being much more heritable than the in-lever. Both are also genetically positively correlated (r = 0.46, SI Fig. 1), and their ratio constitutes the mechanical advantage of the lever system for the superficial masster (M.A. = IL/OL), which directly impacts the proportion of muscular force that can be transmitted to the incisors. Despite the out-lever being the most heritable trait in our study, it appears that the heritability and additive genetic variance of the mechanical advantage is actually restricted by the lower additive genetic variance of the in-lever, combined with the positive additive genetic correlation between the in-lever and out-lever (Fig. 3C, SI Fig. 1). Therefore, it may be that the additive genetic vairance of complex traits such as bite force or mechanical advantage, which depend on numerous inter-correlated other traits, may actually not behave additively. Such an explanation for low heritability of performance or fitness-related traits is not exclusive of other effects, but would require formal testing in the future.

Akin to the model of Schluter (1996), the region of lever arm additive genetic covariation that is compatible with relatively invariant bite forces may form a line of least resistance along which craniofacial morphology may evolve even under stabilizing selection for bite force (Fig 3). This selection on bite force should also produce pleiotropy and/or linkage disequilibrium between morphological traits mechanically related to bite force. That is because in stable conditions the additive genetic variance-covariance pattern will evolve to match the stabilizing selection pattern (Cheverud 1984, 1996). In the case of a many-to-one relationship, variable combinations of traits can be inherited together, as long as they imply performance close to the optimum (Cheverud 1996). This could explain that some genetic covarition patterns of lever-arms or shape do not relate to a major component of mechanical performance variation (Fig. 1, Fig. 3B, SI Fig 1).

Modularity and many-to-one mapping can serve to maintain standing genetic variance without impacting the selection process on the system in normal conditions. The system is therefore not doomed to the reduction of additive genetic variation, and evolution of performance would still be possible if changes occur in the pleiotropic / linkage relationships underlying genetic integration within modules or traits (such as lever arms). Causes of pleiotropic breakdown can be multiple, and genetic correlations have been shown experimentally to be modified under selection, sometimes in opposite directions to the original or expected genetic correlation (Sikkink et al. 2015). It is therefore likely that stabilizing selection would favor modularity and the many-to-one relationship in normal conditions, while directional selection may disrupt these patterns in changing environments.

Fisher’s theorem and the non-expression of GxE interactions (see previous section) could explain the non-detectability of additive genetic variance in bite force in our study. In addition to these phenomena, we propose that non-additive mechanical relationships between components (or functional modules) of a complex performance trait (Fig. 3), provide a non-exclusive and complementary explanation for the reduced heritability of fitness-related and performance traits under stable conditions.

### Implications for the evolution of performance at the interspecific scale

The most heritable morphological traits are size and the out-lever lengths of the mandible, both at the incisor and molar, while the most evolvable trait is the mechanical advantage of the temporal muscle. These traits broadly correspond to a general lengthening (incisor out-lever) or heightening (molar out-lever) of the mandible, and changes in the condyloid and coronoid process (temporal mechanical advantage). These changes are also seen when looking at the genetically heritable shape variation. They can have functional consequences, by modifying the mechanical advantage of the different masticatory muscles, and thereby impacting the amount of force produced and transmitted to food, and the speed of the jaw closure (e.g. Hiiemae 1971, Hiiemae & Houston 1971, Satoh 1997). Comparative morpho-functional studies at the interspecific level in rodents have often noted changes in the length and height of the mandible, as well as in the coronoid and condyloid processes, associated with changes in muscular insertions and tooth morphology (e.g. Michaux et al. 2007, Hautier et al. 2012, Maestri et al. 2016). These trends were linked to ecological (notably dietary) aspects at different scales, within (Ctenohystrica, Hautier et al. 2012; Sigmodontinae, Maestri et al. 2016) and between (Samuels 2009) various groups of rodents. Such morphological changes have been repeatedly selected during over rodent evolution. Our data suggest that size, as well as the lengthening and heightening of the mandible, associated with lever arm changes in the masticatory apparatus, are highly evolvable characters, which may constitute one of the adaptive pathways for rodents to modify their diets or mode of life.

## Conclusion

Here, we report on the finding that bite force, a key measure of performance for craniofacial morphology, is not heritable. This striking result shows that the evolvability of performance traits such as bite force can be complex. We present a hypothesis for the structure of additive genetic variation in the morphological determinants of bite force that is compatible with our data. This hypothesis reveals that an apparent lack of heritability for such a trait within a population does not necessarily mean that the performance trait is not evolvable (Fig. 3B-C). When there are multiple morphological determinants of performance that covary and have a many-to-one, rather than additive, relationship to the functional output, the result is latent genetic variation in performance. This latent variation may be revealed under conditions in which the performance optimum is altered due to environmental change (e.g. in a speciation context), which breaks down genetic correlations between morphological traits or modules involved in producing the performance output.

## Supporting information

Combined SI Text, Figures and Table

