## Supplementary material for "Bite force performance from wild derived mice has undetectable heritability despite having heritable morphological components": Combined SI Text, Figures and Table

### Supplementary Information

**SI Text. Results of multivariate heritability estimates.** Multivariate heritability estimates were of 0.29 for the cranium and 0.32 for the mandible. The latter is in the range of the average univariate heritability estimates for functional traits (bite force excluded) which is 0.31-0.37 for the mixed models and regression estimates, respectively (Fig. 2, SI Table 2). The heritability of cranial and mandibular morphology are therefore confirmed in the multivariate context of shape. However, it should be kept in mind that the assumption of isotropy of matrices underlying the heritability estimator used here was not tested (Monteiro et al. 2002, Klingenberg & Monteiro 2005). This does not allow to draw hypotheses on the potential trajectory that would be followed by skull shape under selection. Still, heritability values for mandible and cranium morphology do match with the expected heritability for morphological characters (Hoffman et al. 2016), and with the average univariate heritability of morphological traits found in the present study (Fig. 2O, SI Table 2).

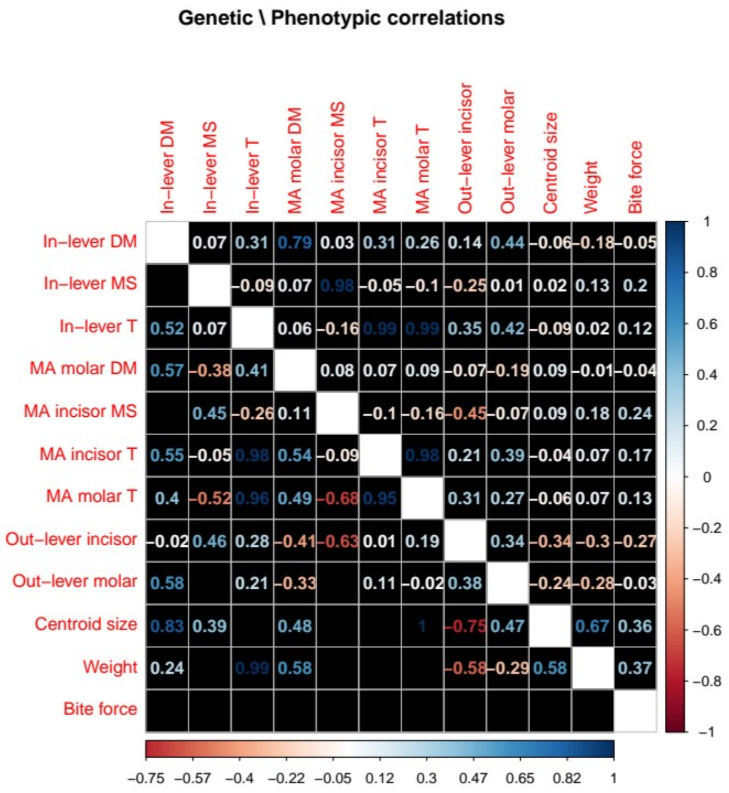


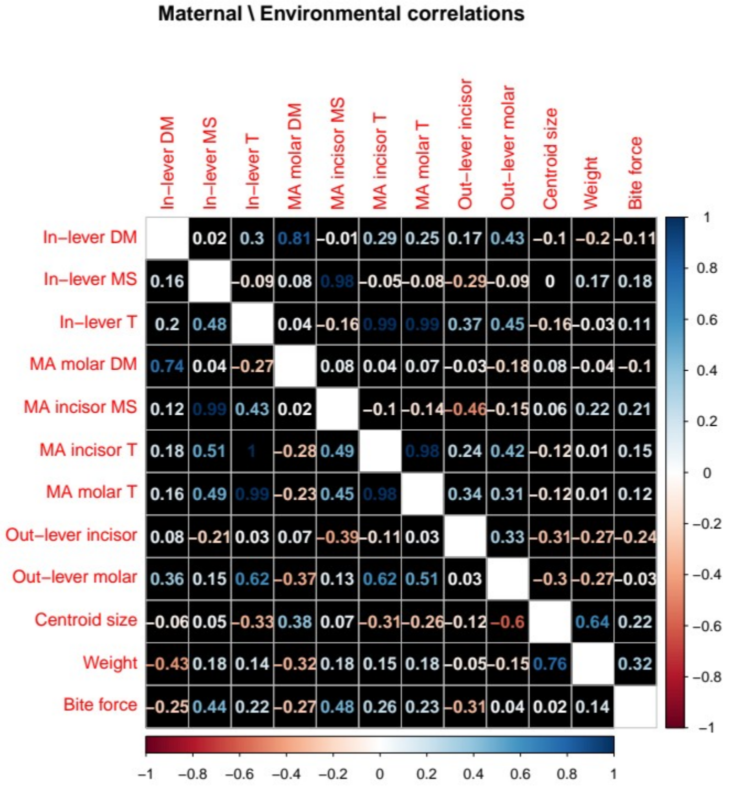


**Supplementary Figure 1.**

Correlation plots of morpho-functional trait data. The upper panel shows phenotypic correlations (upper triangle) and additive genetic (sire) correlations (lower triangle). Note that the latter has missing values (empty black squares). The lower panel shows the residual correlations (upper triangle) and dam correlations (lower triangle).

**SI Figure 2.** Sexual dimorphism in morpho-functional variables. White boxes represent females and gray boxes males. Abbreviations: DM, Deep Masseter; MS, Superficial Masseter; T, Temporal; M.A., Mechanical Advantage.

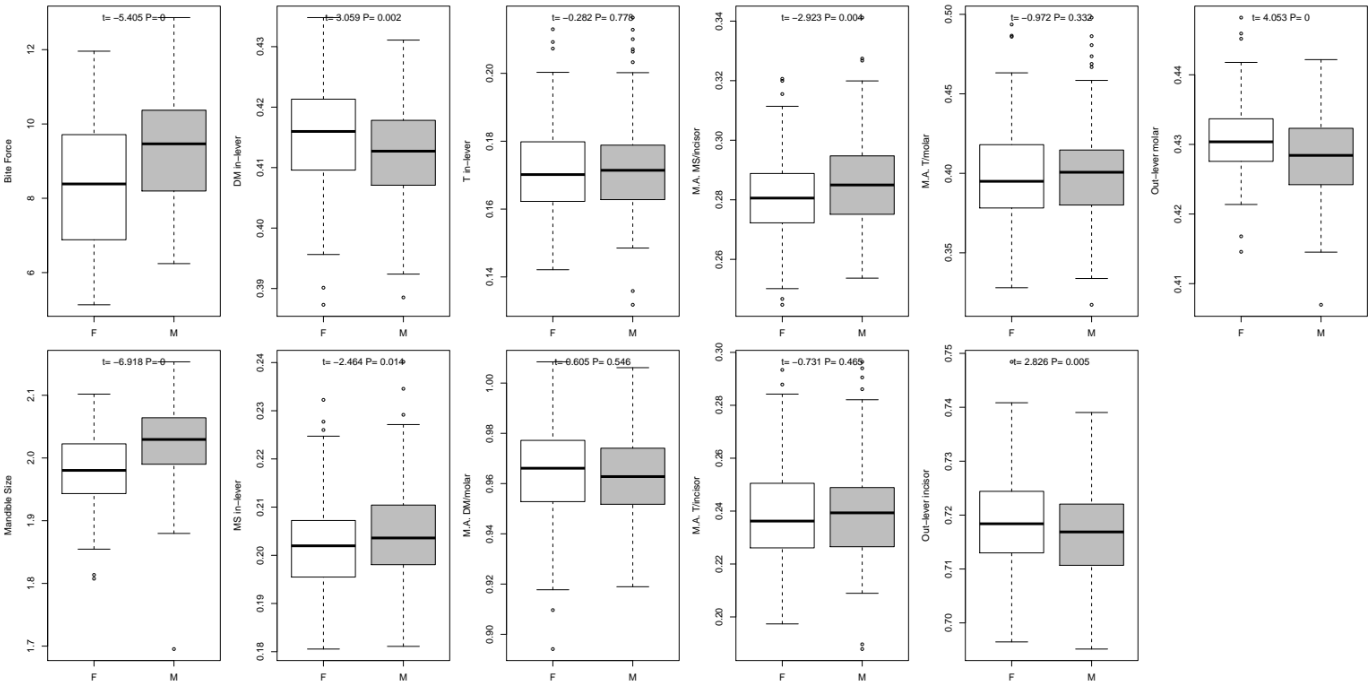

**SI Table 1.** Results of the quantitative genetic analyses ran in this study for traits correlated with bite force. Abbreviations : Cent. Size. : Centroid size ; Mass. Sup : Superficial masseter muscle ; Inc. : Incisor ; MA : Mechanical advantage.

| | Sire | Dam | Residual | $h^2$ (regression) | $h^2$ (mixed model) | Evolvability ($I_A$) |
|---|---|---|---|---|---|---|
| Bite force | 0 | 1,43 | 1,40 | 0 | 0 | 0 |
| Cent. Size mandible | 1,85E-04 | 1,22E-03 | 1,90E-03 | 0,43 | 2,25E-01 | 3,702836e-04 |
| Cent. Size cranium | 3,88E-04 | 2,12E-03 | 3,75E-03 | 0,25 | 2,48E-01 | 4,623796e-04 |
| In-lever Mass. Sup. | 9,00E-07 | 2,11E-05 | 7,12E-05 | 0,15 | 4,01E-02 | 1,839904e-05 |
| MA Mass. Sup./Inc. | 2,30E-06 | 4,61E-05 | 1,64E-04 | 0,14 | 4,38E-02 | 3,287980e-05 |
| MA Temp./inc | 2,27E-05 | 5,79E-05 | 2,37E-04 | 0,4 | 2,86E-01 | 3.802702e-04 |
| Out-lever incisor | 1,75E-05 | 1,03E-05 | 4,13E-05 | 0,67 | 1,01E+00 | 9,746540e-05 |
| Out-lever molar | 4,23E-06 | 4,66E-06 | 2,10E-05 | 0,53 | 5,57E-01 | 3,942059e-05 |
